# Prediction of liquid-liquid phase separation proteins using machine learning

**DOI:** 10.1101/842336

**Authors:** Tanlin Sun, Qian Li, Youjun Xu, Zhuqing Zhang, Luhua Lai, Jianfeng Pei

## Abstract

The liquid-liquid phase separation (LLPS) of bio-molecules in cell underpins the formation of membraneless organelles, which are the condensates of protein, nucleic acid, or both, and play critical roles in cellular functions. The dysregulation of LLPS might be implicated in a number of diseases. Although the LLPS of biomolecules has been investigated intensively in recent years, the knowledge of the prevalence and distribution of phase separation proteins (PSPs) is still lag behind. Development of computational methods to predict PSPs is therefore of great importance for comprehensive understanding of the biological function of LLPS. Here, a sequence-based prediction tool using machine learning for LLPS proteins (PSPredictor) was developed. Our model can achieve a maximum 10-CV accuracy of 96.03%, and performs much better in identifying new PSPs than reported PSP prediction tools. As far as we know, this is the first attempt to make a direct and more general prediction on LLPS proteins only based on sequence information.

## Introduction

The efficiency and precision of bio-chemical reactions in eukaryotic cells are much attributed to the spatial temporal compartmentalization of bio-molecules in cytoplasm[1]. Compartments have traditionally been thought as membrane-bound organelles such as endoplasmic reticulum. Another category, membraneless organelles (also called bio-molecular condensates) which were observed many years ago, have recently been recognized to compartmentalize cellular space through liquid-liquid phase separation (LLPS)[2,3]. More and more studies suggested that many cellular metabolic processes are regulated by LLPS, so are some intractable diseases such as ALS (amyotrophic lateral sclerosis)[4] and AD (Alzheimer disease)[5]. A number of proteins have been verified to be able to form liquid-like membraneless assemblies in cells and *in vitro* [3,6–9]. Meanwhile, studies indicated that RNA can regulate the LLPS of proteins and influence the formation of bio-molecular condensates[10]. Fundamentally, the reversible multi-valent interactions between bio-molecules drive LLPS process, which can be occurred between multiple folded domains or be mediated by intrinsically disordered proteins (IDPs). Generally, phase separation related proteins can be categorized as scaffolds which drive LLPS, and clients, which participate into the condensates formed by scaffolds[11,12]. Although tremendous progress has been made in understanding protein LLPS, the knowledge of prevalence and distribution of phase separation proteins (PSPs) is still lacking. Development of computational methods to predict PSPs is therefore of great importance for the knowledge expansion of LLPS.

In a recent review, Vernon et al. summarized a range of first-generation PSP prediction tools[13]. Each of them is based on specific protein features which are deemed as the driving force of LLPS. Specifically, PScore is based on the expected number of long-range, planar sp2 pi-pi contacts[14]; DDX4-like is based on sequence composition and residue spacing similarity to DDX4[15]; PLAAC is based on prion-like domains[16]; LARKS is based on low-complexity aromatic-rich kinked segments[17]; R+Y is based on the proportion of arginine and tyrosine and the features of FET family proteins[18]; and CatGranule is based on statistical analysis of amino acids composition responsible for granule forming[19] Recently, Orlando et al. predicted FUS-LIKE PSPs by Hidden Markov Model (HMM), and the model considered prion-like domains (PLD), disordered region, arginine rich domain, RNA recognition motifs (RRM) and other features [20]. Their tool (PSPer) has been successfully used to predict the phase separation ability for 22 experimentally studied FUS-LIKE proteins[18]. However, all these methods were based on small samples and specific features, which hampered the scopes of their applications. Large data-based prediction tools with more general application scopes are in urgent need.

Machine learning is one of the most powerful methods for prediction of protein functions. Various aspects of protein features can be integrated into descriptors or vectors by using physical, chemical properties of residues or context of sequences to train a prediction model. Development of PSP prediction tools by using machine learning have been hampered due to lacking the collection of experimentally determined PSPs data. Recently, a number of PSP databases have been published [21–23], they set the ground for the birth of more general PSP prediction tools. In particular, Li et al. curated a LLPS database (LLPSDB)[24] by extracting LLPS experimental results from literatures. Each entry in the database contained the information of whether the protein (alone or with DNA/RNA or with other proteins) phase separated under a specific experimental condition *in vitro*. In this work, we developed a sequence-based machine learning PSP prediction tool (PSPredictor) based on the dataset in this database. As far as we know, this is the first attempt to make a direct and more general prediction on proteins undergoing LLPS based on sequence information. Our model achieved a 10-CV training accuracy of 96.03% and a 90.90% prediction accuracy on an external test set containing 88 proteins, and performed much better in identifying new PSPs than other reported PSP prediction tools so far. The limitations of our model, as well as the problems to be solved further are also addressed.

## Results

### Dataset construction for PSP prediction

In this paper, we defined PSPs as proteins that can undergo LLPS on their own or with DNA/RNA. For model training, we used PSP sequences in LLPSDB beta database that were experimentally validated to undergo LLPS on their own (single protein system), or with DNA/RNA, as positive training dataset (dataset A1, 284 protein sequences). All of them contained IDRs (or are IDPs themselves), without post-translational modification. As LLPS is driven by multivalent interactions between multiple rigid domains or disordered ones, we collected single-domain proteins in the PDB databank with full length three-dimensional (3D)-structures solved as the negative training dataset (B1), which contains 5258 protein sequences.

### Development of the PSP prediction tool – PSPredictor

In order to find which model performed the best, three categories of variables, including ratio between positive and negative samples(1:1,1:2 and 1:5), as well as selected protein coding methods (word2vec, w2v for abbreviation; and Li’s method, LQL for abbreviation), and seven machine learning algorithms (Supported Vector Machine (SVM), Random Forest(RF), Logistic Regression(LR), Decision Tree (DT), Gradient Boosting Decision Tre e(GBDT). Naive Bayesian (NB)) (see the METHOD DETAILS) were systematically combined to build a total of 42 (3 x 2 x 7) models. By considering the statistical indexes of accuracy, F1, precision, recall specificity and MCC (≥90%), the best model (Model 1) was selected which is w2v coded, trained by GBDT, and the ratio between positive and negative samples is 1:1. It achieved a 10-CV training accuracy of 96.03%±1.30% (the training statistical index values are shown in Table 1). Because the negative samples were randomly selected from dataset B1, we repeated the model training independently three times, and all the training results can be found in Table S1, S2, and S3. More details of the construction of training datasets, protein coding methods, machine learning algorithms, as well as the definition of statistical indexes can be found in METHOD DETAILS.

**Table 1.**
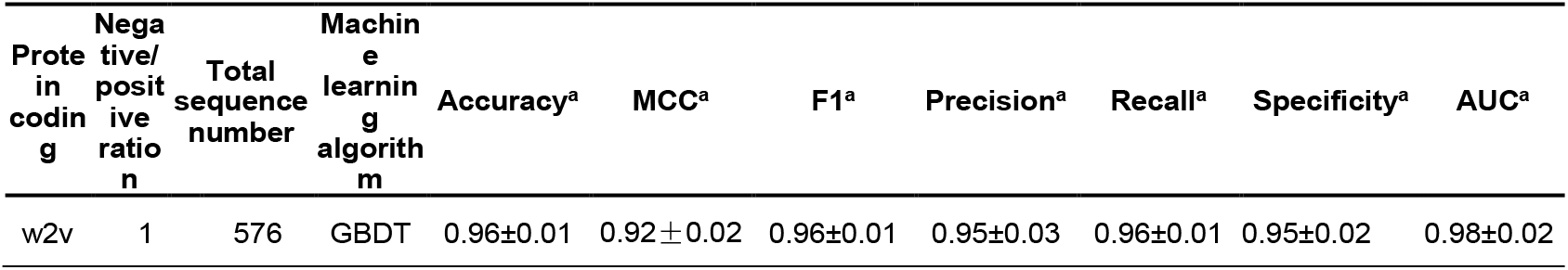
The evaluation of the best model (Model 1) for PSPs prediction

We selected the first trained best model (Model 1) from the three repeats to predict the external dataset. The positive dataset F1+ contains 88 newly added protein sequences (NAPS, see the METHOD DETAILS) in LLPSDB release, comparing with LLPSDB beta. Although dataset F1+ shares low sequence similarity (only 20% proteins in dataset F1+ have ≥ 30% sequence similarity with those in dataset A1) or functional similarity (e.g. RNA Recognition Motif (RRM) is enriched in dataset A1, while no proteins in dataset F1+ have RRM) with dataset A1, the prediction accuracy is 90.90%. Meanwhile, we used the dataset B1 excluding the sequences in the negative training dataset as external negative test sett). The prediction accuracy is 93.20%.

We also tested two first generation PSP prediction tools, PScore and CatGranule, which performed best among the 7 first-generation methods[13] and PSPer[20], on dataset F1+. The relationships between percent recall and total percentage of whole proteins accepted at given thresholds, for PScore, CatGranule, PSPer and our Model 1, are shown in Figure 1. Obviously, our Model 1 is superior to other models in dataset F1+ prediction.

**Figure 1.**
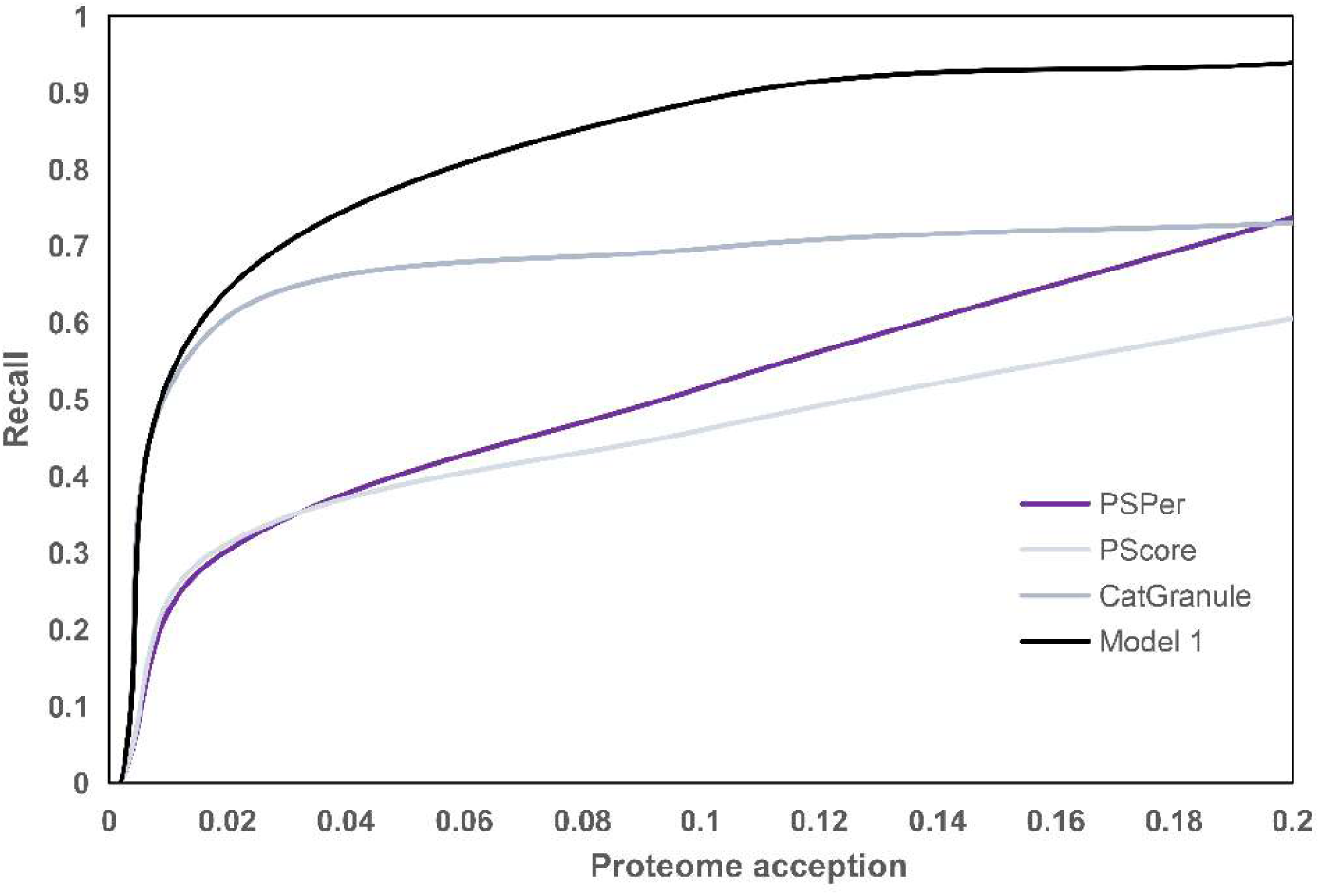
Relationship between percent recall and total percentage of human proteins accepted at given thresholds, for Model 1 and three best first generation prediction tools.

We combined dataset A1 and F1+ together as a full positive training dataset (A2), and used the same negative dataset and parameters of Model 1 to train a model, that is, PSPredictor as our final PSP prediction model.

### Analysis of scaffolds, regulators, clients and granule forming proteins

Proteins involved in LLPS can be categorized as scaffold proteins (scaffolds) and client proteins (clients)[11,12]. Scaffolds are defined as the drivers of LLPS while clients have been discovered to co-occurrence with scaffolds in experimental conditions, but lack of evidence of whether they can undergo LLPS or not independently. Recently, Ning et al. published a database, DrLLPS[25], in which LLPS-related proteins were further categorized as scaffolds, regulator proteins(regulators, responsible for regulating the LLPS behaviors of scaffolds, by various mechanism such as post-translational modification) and clients. However, these proteins sometimes overlapped, which means individual protein may act as scaffold, regulator or client depending on various distinct systems[26]. To give an estimation of real PSPs, which means proteins can undergo LLPS on their own or with DNA/RNA, by our scope of definition, we first analyzed the relationships between the positive training dataset of PSPredictor (dataset A2) and three categories proteins in DrLLPS. It is well accepted that IDR is one of the critical driving force of LLPS. The average disorderness of scaffolds is the highest among the three categories (0.24±0.21,see METHOD DETAILS), and in dataset A2 the average disorderness of scaffolds is even higher (0.3±0.17). Furthermore, dataset A2 share the highest sequence similarity with scaffolds than the other categories (about 70% sequences share more than 80% sequences similarity with scaffolds). At a high threshold (1.8%), around 50% of scaffolds are predicted as PSPs by PSPredictor, while only 10.54% regulators and 8% clients are predicted as PSPs. Also, the proportions of PSPs predicted by PSPredictor are higher than that of PScore (Figure 2).

**Figure 2.**
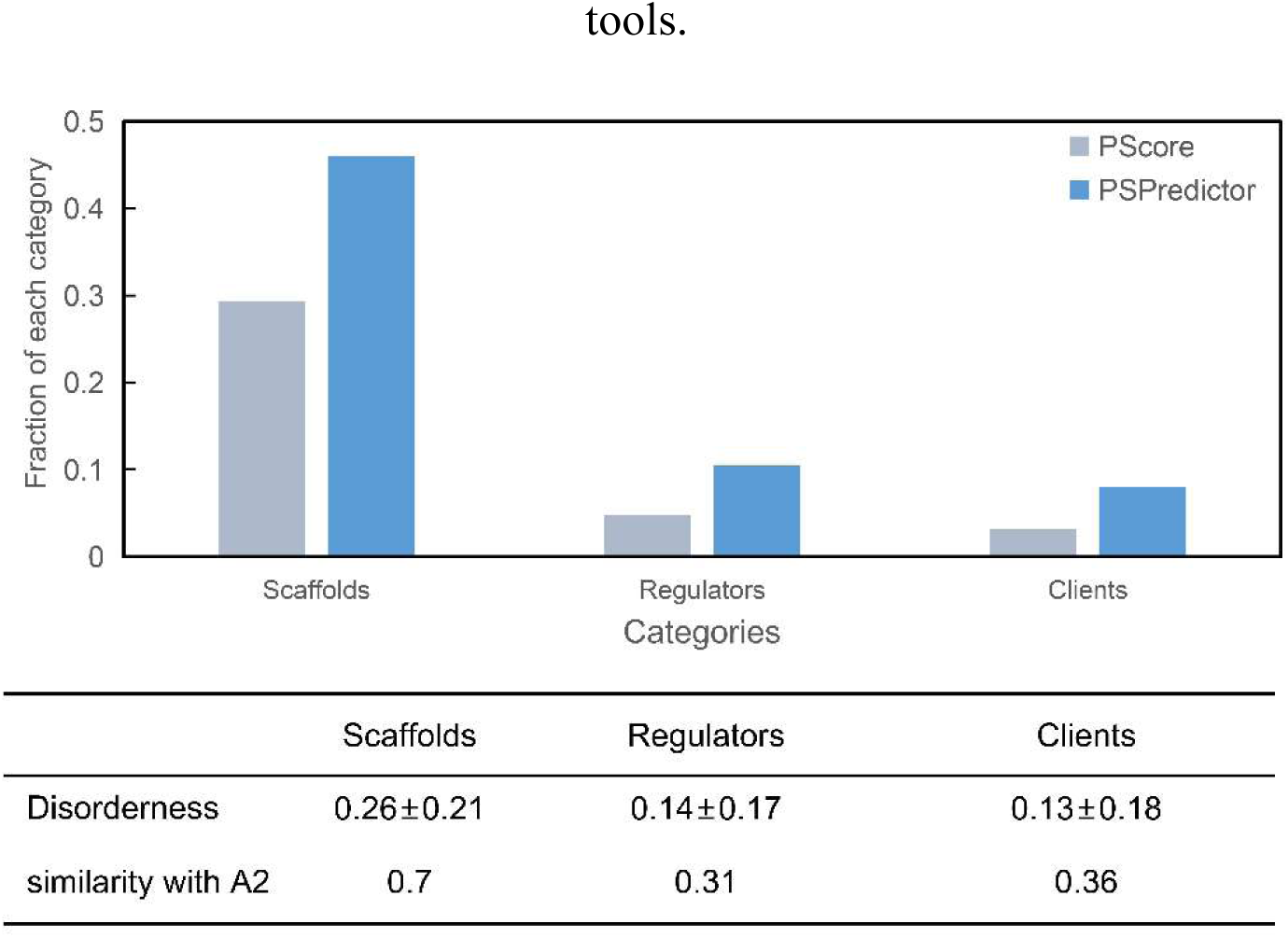
Fraction of proteins in each categories (scaffolds, regulators or clients) predicted as PSPs by PSPredictor or PScore. And the disorderness of each category, the proportion of sequences in A2 that shared ≥ 80% similarity with each category are showed below.

Some proteins are the core components of stress granules or P-body condensates, but whether they could undergo LLPS on their own were not known. Recently, Youn et al. published a comprehensive database containing the proteins related to the formation of stress granules and P-body condensates. Protein was each assigned to tier 1-4 according to the degree of confidence in evidence supporting protein localization to stress granules or P-body (4385 protein sequences)[21]. We predicted these data by using PSPredictor and compared the results with those of the reported PSP prediction tools. The prediction result for each tier is shown in Figure 3A. We also calculated the number of overlapped PSPs predicted between any two tools and the overlapping matrix is shown in Figure 3B. For all prediction tools, the proportions of predicted PSPs ranked in the order of tier 1 > tier 2 > tier 3 > tier 4, which is in accordance with the degree of confidence in evidences defined by Youn et al.. PSPredictor predicted more PSPs than other tools, and sharing more overlapped predicted PSPs with other tools. For all the proteins in the database reported by Youn et al., PSPredictor prediction indicates that around 9.9% of the proteins in stress granules or P-body may undergo LLPS spontaneously, while others gave an estimation of 3-4%.

**Figure 3.**
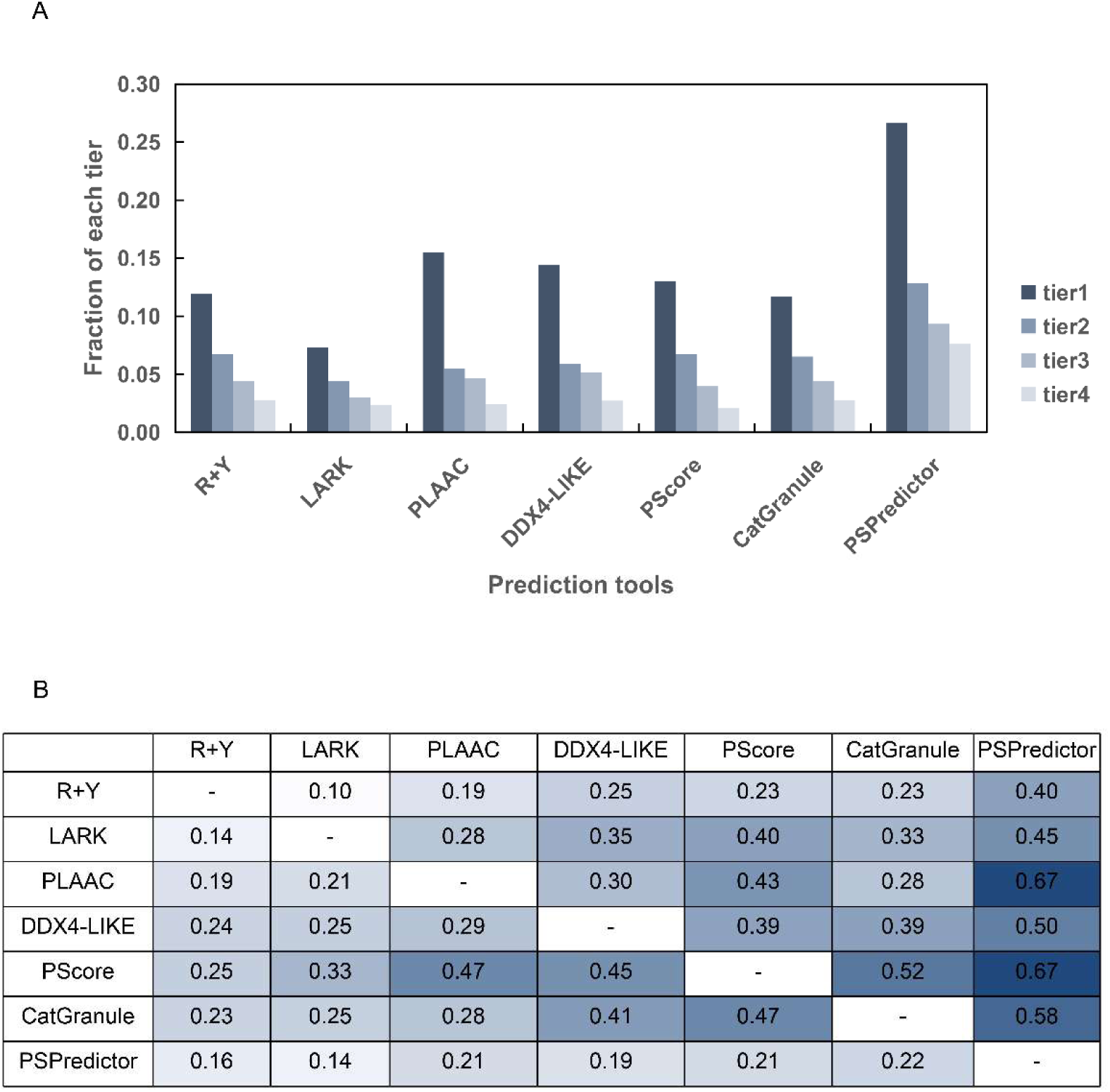
(A) Fraction of proteins in each tier group predicted as PSPs by first generation prediction tools and PSPredictor. (B) The number of overlap predicted PSPs by each two prediction tools.

These results re-emphasize that there are only a small proportion of proteins spontaneously involved in the formation of condensates[20,26], even for the scaffolds, a large proportion of which might participate in multi-component LLPS environments. While some proteins temporarily categorized as regulators and clients might also undergo LLPS on their own in other systems, which remain to be validated in the future.

### Scanning the human genome for potential PSP

Human proteins in the top 1.8% (high confidential threshold) scores of the whole human proteome predicted by PSPredictor are regarded as consolidated PSPs. GO terms enrichment analysis on these predicted PSPs was performed and the terms with EASE score <0.1 are showed in Table S4 (see also the METHOD DETAILS). Most of the GO terms that were identified by first-generation PSP prediction tools [13], are also enriched in our predicted consolidated PSPs, such as cytoplasmic stress granule, ribonucleoprotein complex, etc. Comparing with first-generation tools, nucleus associated PSPs are more likely to be identified by PSPredictor.

When clustered GO terms with similar biological context at a significant threshold (1.8%), 12 clusters with high enrichment scores stand out (Table S5): For proteins in the *collagen trimer, endoplasmic reticulum lumen, and extracellular matrix related proteins* cluster, they provide structural support, biochemical or biomechanical cues for cells or tissues. Phase separation might be involved in the activities of those proteins, and phase separation related techniques had been recruited to create biomimetic artificial nano materials for cells or tissues culturing[27,28]. Dentin sialophosphoprotein 1, enamelin, etc. are such proteins predicted as PSPs by PSPredictor, but had rarely been studied; For proteins in the *nBAF complex, chromatin remodeling and SWI/SNF complex related proteins* cluster, they are responsible for chromatin remodeling and gene expression regulation. LLPS had been reported to play critical roles in these two processes[29,30]. SWI/SNF were reported to be enriched at NLIs, which are a novel type of nuclear PIP2-enriched structures containing proteins, lipids and RNA - the typical components of phase separated nuclear compartment proteins[31]. SMARCC2 and SMARCA2, which are the components of SWI/SNF, were predicted as PSPs by PSPredictor. The LLPS behaviors of these proteins have not been much studied before. This indicates PSPredictor can provide new clues for the functional studies of newly predicted PSPs.

## Discussion

### The machine learning algorithm and sequence encoding methods used in PSP prediction

Most of our best models were obtained by training using GBDT, one of the most powerful and widely used supervised machine learning algorithms. It is the integration of gradient boosting and decision trees and has the ability for both linear and nonlinear data classification, regression and prediction. GBDT can generalize and combine weak learners into a single strong learner and has produced good results in biologic data mining, comparing to other machine learning algorithms [32–36]. Our research is another successful application of GBDT in biologic field.

W2v is one of the natural language processing techniques by which words are embedded in vectors through the training of contexts. They can also embedded residues, proteins and chemicals into vectors as the input for model training without artificial design of features or expert knowledge. It had been used to predict HLA binding proteins, antimicrobial peptides and drug targets [36–39]. Our current research validates its capacity in PSP prediction.

### Datasets used for training PSPredictor

We used the PSPs in LLPSDB as our positive dataset, which includes IDPs as well as proteins with IDRs; while our negative dataset contains single-domain proteins with full length 3D structures solved. Therefore one might doubt that our model may become a kind of IDR prediction model. To rule out this possibility, we compared the prediction scores for proteins in the dataset A1 and IDPs from Disprot. There were significant differences (p<0.01) between the scores of two protein groups, and only 31% proteins in the IDP dataset were predicted as PSPs (see METHOD DETAILS). These results indicate that PSPredictor is not an IDR prediction model. Since the positive dataset did not include proteins without evidence suggesting that they could undergo LLPS on their own, or in single protein system, or became PSPs only after post-transcriptional modifications, PSPredictor may not be good at in those proteins prediction.

### Perspectives for future PSP prediction

We have shown that with the data collected in LLPSDB, it is possible to develop universal PSP prediction model that is not restricted by a few specific protein domains. With limited experimental data available before, most of the first-generation PSP prediction tools depend on specific domains. Along with more accumulated PSP data, we expect that predictive tools like PSPredictor can cover more PSP space with high prediction accuracy. Other data-demanding algorithms can also be employed in the future. Currently, PSPredictor as well as the first generation prediction tools can be used to predict driver proteins, while client proteins may need specialized prediction tools or whether generalized tools can be developed need further investigation.

Generally speaking, every protein can undergo LLPS under correct conditions. In our positive dataset, we only included PSPs that form LLPS under near physiological conditions without considering their differences. With more data available, it may be possible to include experimental conditions in the training process, so that PSP can be predicted with experimental conditions like temperature, salt, pH and crowding agents. The other challenge is to predict PSP mutants that destroy LLPS process. Due to the high level of sequence similarity, it is difficult for sequence based prediction tools to differentiate them. More features may be required for this kind of prediction.

## Supporting information

all supplementary files

## Acknowledgments

We thank the suggestions and helps from Dr. Weilin Zhang, Dr. Hao Liang and associated professor Changsheng Zhang. This work was supported in part by the Ministry of Science and Technology of China (2016YFA0502300, 2015CB910300) and the National Natural Science Foundation of China (21673010, 21633001).

## Author Contributions

T.L.S. performed the model training and data analysis. Q.L. and Z.Q.Z. built and curated the LLPSDB. Y.J.X built the computation server (in construction). J.F.P., Z.Q.Z. and L.H.L. conceived the project. T.L.S., J.F.P., Z.Q.Z. and L.H.L wrote the paper.

## METHOD DETAILS

### Training and Test dataset construction

For the LLPSDB, we defined a *condition associated* entry (condition, for short) as a protein sequence and its LLPS behavior (positive, if it could undergo LLPS; negative, if it could not undergo LLPS) under a specific condition(pH, temperature and salt concentration, etc.); we defined a *sequence* entry as a protein sequence and its PSP behavior (if the protein sequence could undergo LLPS in at least one condition, it was believed to be a positive sequence); we defined a *protein* entry as a protein (without considering its mutations) and its LLPS behavior. The number of overall condition associated entries, sequence entries and protein entries were illustrated in Figure S1. For all the PSPs, only the simple coacervation entries were used, which means that the system only includes intrinsically disordered region (IDR)-containing protein undergoing LLPS in solution by itself or with RNA/DNA.

For candidate negative dataset, we downloaded sequences from the PDB [40] and selected those with 3D-structures covering the full-length sequences. There were 14778 of such protein sequences. We removed redundancy so that no two sequences in the dataset were allowed to share sequence similarity ≥ 50%. A total of 8625 protein sequences were obtained. Then, Hmmer3 was used for domains prediction, and single domain protein sequences were selected as Negative Dataset B1 (5258 sequences). Negative dataset with protein samples from LLPSDB beta was also constructed for illustrating question 4 in Discussion section.

#### Positive Dataset A1

Protein sequences in LLPSDB beta that could undergo LLPS on their own or with DNA/RNA. We deleted the sequences whose 1) length were lower than 50, 2) who contain B, U, X that could not be coded by our coding methods, 3) who are regulated by PTM, 4) no evidence indicated that it can undergo LLPS in single protein system. A total of 284 protein sequences were selected.

#### Negative Dataset B1

Protein sequences which were unlikely to undergo LLPS. A total of 5258 protein sequences were selected. When dataset B1 was used as training, the equal, 2-fold, or 5-fold number (as compared to the number of positive samples) of protein sequences of dataset A1 were selected randomly as negative samples. When dataset B1 was used as test set, since only a small part of it was used as training samples, they were not removed from dataset B1.

#### Positive Dataset F1+

The current LLPSDB release has more entries than LLPSDB beta. We used the newly added protein sequences (NAPS) in the LLPSDB release (other than LLPSDB beta) that could undergo LLPS to construct the test datasets. A total of 88 protein sequences were selected.

#### Positive Dataset A2

A combination of dataset A1 and F1+. A total of 372 protein sequences were obtained.

#### Negative Dataset B2

Protein sequences from LLPSDB beta that without evidences to undergo LLPS. A total of 105 sequences were obtained. Since the number of sequences in B2 was much less than sequences in A1, the equal, 1.5-fold, 2-fold number (as compared to the number of negative samples) of protein sequences of dataset B2 from dataset A1, or the whole set of A1 were selected randomly as positive samples, when dataset B2 was used as training.

#### Youn’s dataset

This dataset contained 4385 stress granule or P-body related proteins, which were assigned to 4 tiers based on the quality and sufficiency of evidences to undergo LLPS. Tier 1 contains 368 proteins, tier 2 contains 475 proteins, tier 3 contains 428 proteins and tier 4 contains 3114 proteins.

#### DrLLPS

This database contained 150 scaffolds, 986 regulators and 8148 clients[25]. They were collected from PubMed.

#### IDP dataset

The IDPs dataset was downloaded from Disport 8.0[41] (https://www.disprot.org/). We removed redundancy so that no two sequences in the dataset were allowed to share sequence similarity ≥ 50%. A total of 1339 protein sequences were remained. When threshold was set to score=0.5, the number of correctly predicted PSPs were calculated for IDP dataset and external test dataset (dataset A1). Since the score distributions were not normal, Mann-Whitney U test was used to compare the score rank between 30 human PSP dataset and the IDPs dataset (SPSS 24.0). The average score for IDP dataset is 0.391, and the average score for external dataset is 0.883, there are significant difference between the two groups (p<0.01).

### Protein coding

Our previous experiences showed that LQL method and w2v were better in protein coding. They could capture important features and performed well in model training and prediction[36].

#### LQL method [42]

The composition of 20 amino acids formed the first 20 dimensions. Then the amino acids were clustered into 3 types of amino acids for each of six physicochemical property (hydrophobicity, polarity, polarizability, solvent accessibility and normalized van der Waals volume). Composition, Transition and Distribution of each type and each amino acid type property was calculated. For example, amino acids were clustered into polar, neutral and hydrophobic for hydrophobicity property. For ‘Composition’, 3 dimensions were calculated: the percentage of polar, neutral and hydrophobic; For ‘Transition’, 3 dimensions were calculated: the percentage of polar transferred to neutral, neutral transferred to hydrophobicity and hydrophobicity transferred to polar; For ‘Distribution’, 5 dimensions were calculated for each of the 3 types of amino acids: the location percentage of the first, 25%, 50%, 75% of that type. A vector of total 146 dimensions were calculated for LQL method.

#### Word2vec(w2v)

W2v[43] is the name of a series of models that are trained to produce word embedding vectors. Two models are popular: continuous bag-of-words (CBOW) and continuous skip-gram. The former predicts current word from a window surrounding the context words, while the latter uses the current word to predict the surrounding window of context words. Hierarchical softmax and negative sampling are the two main training methods. The former uses a Huffman tree to maximize the conditional log-likelihood, while the latter minimize the log-likelihood of sampled negative instances. In this work, Skip-gram model with window size 8, and hierarchical softmax were recruited. We downloaded the whole protein sequences from swiss-prot, and broke the original sequences into 3 residue-length window overlapped kmers. The dimension was set to 200. We used w2v program in genism python NLP package [44](https://radimrehurek.com/gensim/) to train and compute the embedding vectors. Once obtaining the embedding vectors, we added the 3-gram vectors in each sequences to form the sequence vectors.

### Machine learning

Machine learning models: Supported Vector Machine (SVM), Random Forest (RF), Logistic Regression (LR), Decision Tree (DT), Gradient Boosting Decision Tree (GBDT). Naive Bayesian (NB) in scikit-learn (https://github.com/scikit-learn/scikit-learn) were recruited. The parameters for SVM, RF, LR, GBDT and KNN were adjusted by grid search, while the parameters of DT and NB were used as default. For SVM, Radial Basis Function (rbf) was used as the kernel function and grid search was done to C and gamma. For RF, grid search was done to the number of estimators. For LR, grid search was done to C. For GBDT, grid search was done to learning rate and the number of estimators. For KNN, when weights was set to ‘uniform’, grid search was only done to the number of neighbors; when weights was set to ‘distance’, grid search was done to the number of neighbors and p(distance metric).

Machine learning models were evaluated by Accuracy, Area under curve (AUC), Matthews’s correlation coefficient (MCC), F1, Recall, and Precision.

TP (True positive): The number of proteins predicted as PSPs and they were real PSPs.

TN (True negative): The number of proteins predicted as none-PSPs and they were theoretically unlikely to undergo LLPS.

FP (False positive): The number of proteins predicted as PSPs but in fact they were theoretically unlikely to undergo LLPS.

FN (False negative): The number of proteins predicted as none-PSPs but were actually PSPs.

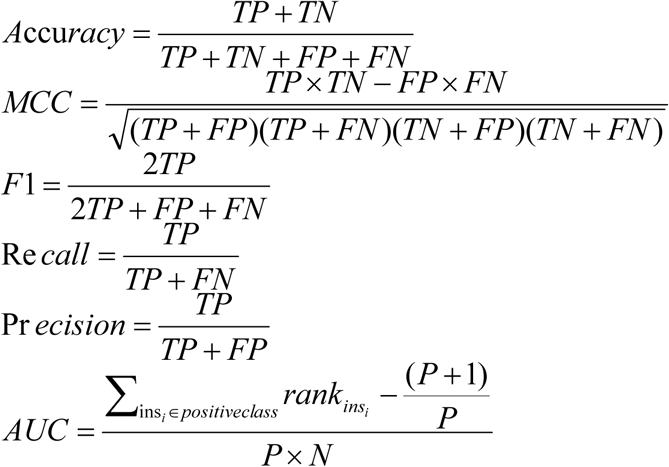

In AUC, rank_ins_ means the rank of a sample sorted by probability scores. P means the total number of positive samples, N means the total number of negative samples.

For our best model (Model 1), the GBDT model parameters were: n_estimators=200, learning rate=0.1, loss=deviance, alpha=0.9, max_features= None, max_depth=None, subsample=1, min_samples_split=2, min_samples_leaf=1, max_leaf_nodes=None, min_impurity_split=1e-7, presort=FALSE.

Considering that the size of B1 is much larger than A1, we repeated the training process for three times, each time the negative training samples were randomly selected from B1. In the main text, we only presented one set of 10-CV training results and the details of all sets of 10-CV training results can be found in Table S2 and Table S3.

### PSP/Non-PSP separation threshold setting

Considering the uncertainty of the negative dataset and that most of the first generation tools only returns prediction scores, it is necessary to set a reasonable PSP/Non-PSP separation threshold Previous tools set this threshold by using the whole proteome as a background, for example, top 1.8%[13], 2%[21] and 20%[13], scored proteins in whole proteome were regarded as PSPs and were regarded as Non-PSPs if not. In this work, for the positive dataset F1+, we used a recall-receive curve to analyze the recalls under different threshold of top n percent of the proteome scores (n was between 0 and 25%). For Youn’s dataset, we set it to 2%, which was the same as Youn’s et al.. For model training accuracy calculation and IDP prediction, score of 0.5 (instead of a background threshold) was used as a direct threshold. For human proteome scan, in order to identify most likely PSPs, 1.8% was set as the threshold.

### Human PSPs GO terms enrichment analysis

Human proteins with scores in the top 1.8% of human proteome were uploaded to DAVID 6.8 (https://david.ncifcrf.gov/)[45]. The enrichment of cellular components GO terms were analyzed in human PSPs, and each of the term got a p-value (EASE score). These yielded GO terms were compared with Vernon et al.’s paper[13], in which enriched GO terms of PSPs predicted by the first-generation tools were presented. Then, for annotation cluster analysis, cellular components, biologic process and molecular function GO terms were analyzed. Geometric mean of EASE scores of those terms (EASE scores < 0.1) with similar biologic meanings were calculated to get the enrichment score for each cluster. Those GO term clusters with high enrichment scores were the most related functions of PSPs predicted by PSPredictor.

### Disorderness calculation

Sequence disorderness proportion was calculated by using MobiDB 3.0[46]. MobiDB could only calculate wild type protein given Uniprot ID. For positive training dataset, where many protein sequences are mutated or truncated, only wild type protein sequences were calculated.

### Sequence redundancy removal

Sequence redundancy removal was conducted so that no two sequence in the dataset were allowed to share sequence similarity equal or larger than a given value. In this study the value was set to 50%. CD-hit[47] git (http://weizhongli-lab.org/cd-hit/) remote package was used to perform these tasks by sequences clustering, the represented sequence in each cluster was selected to form the redundancy removal datasets.

### Sequence comparison

NCBI blast+ 2.2.31[48] remote package was used to perform sequence comparison between two protein sequences.

### Sequence domain annotation

Hmmer 3.1b2[49] (http://hmmer.org/publications.html) remote package was used to perform sequence domain annotation.

## Supplementary Information

Table S1, S2, S3 ALL models training results (repeated three times with different negative samples)

Table S4 Enrichment GO terms of human PSPs predicted by PSPredictor

Table S5 GO term clusters of PSPs with similar meaning in biology

Figure S1. The number of overall conditions associated entries, protein sequences and proteins in LLPSDB beta.

