## Supplementary material for "Prediction of liquid-liquid phase separation proteins using machine learning": all supplementary files

Table S1. related to Table 1
ALL models training results_1

| Protein codings | Ratio of negative/positive | Sequence number | Machine Learning algorithms | Accuracy | F1 | Precision | Recall | Specificty | AUC | MCC |
|---|---|---|---|---|---|---|---|---|---|---|
| LQL | 5 | 1740 | DT | 0.9368 | 0.8116 | 0.8169 | 0.8103 | 0.9621 | 0.8862 | 0.7751 |
| w2v | 1 | 580 | DT | 0.9017 | 0.9029 | 0.9011 | 0.9103 | 0.8931 | 0.9017 | 0.8089 |
| w2v | 2 | 870 | DT | 0.9161 | 0.8758 | 0.8657 | 0.8897 | 0.9293 | 0.9095 | 0.8147 |
| w2v | 5 | 1740 | DT | 0.9448 | 0.8322 | 0.8413 | 0.8276 | 0.9683 | 0.8978 | 0.8008 |
| LQL | 1 | 580 | DT | 0.9017 | 0.9030 | 0.8975 | 0.9103 | 0.8931 | 0.9017 | 0.8051 |
| LQL | 2 | 870 | DT | 0.9264 | 0.8912 | 0.8852 | 0.9000 | 0.9397 | 0.9198 | 0.8372 |
| w2v | 5 | 1740 | GBDT | 0.9782 | 0.9337 | 0.9477 | 0.9207 | 0.9897 | 0.9870 | 0.9210 |
| LQL | 5 | 1740 | GBDT | 0.9764 | 0.9267 | 0.9647 | 0.8931 | 0.9931 | 0.9871 | 0.9142 |
| w2v | 1 | 580 | GBDT | 0.9603 | 0.9604 | 0.9599 | 0.9621 | 0.9586 | 0.9869 | 0.9218 |
| w2v | 2 | 870 | KNN | 0.9598 | 0.9372 | 0.9678 | 0.9103 | 0.9845 | 0.9842 | 0.9096 |
| LQL | 1 | 580 | GBDT | 0.9431 | 0.9424 | 0.9488 | 0.9379 | 0.9483 | 0.9866 | 0.8880 |
| w2v | 2 | 870 | GBDT | 0.9586 | 0.9373 | 0.9456 | 0.9310 | 0.9724 | 0.9870 | 0.9075 |
| LQL | 5 | 1740 | KNN | 0.9557 | 0.8490 | 0.9737 | 0.7552 | 0.9959 | 0.9524 | 0.8338 |
| LQL | 2 | 870 | GBDT | 0.9586 | 0.9358 | 0.9684 | 0.9069 | 0.9845 | 0.9876 | 0.9073 |
| w2v | 1 | 580 | SVM | 0.9586 | 0.9587 | 0.9566 | 0.9621 | 0.9552 | 0.9880 | 0.9184 |
| w2v | 5 | 1740 | KNN | 0.9661 | 0.8909 | 0.9455 | 0.8448 | 0.9903 | 0.9788 | 0.8739 |
| LQL | 1 | 580 | KNN | 0.9000 | 0.8934 | 0.9513 | 0.8448 | 0.9552 | 0.9634 | 0.8070 |
| LQL | 2 | 870 | KNN | 0.9184 | 0.8636 | 0.9714 | 0.7793 | 0.9879 | 0.9534 | 0.8174 |
| LQL | 5 | 1740 | LR | 0.9454 | 0.8091 | 0.9530 | 0.7069 | 0.9931 | 0.9812 | 0.7916 |
| w2v | 2 | 870 | SVM | 0.9575 | 0.9324 | 0.9816 | 0.8897 | 0.9914 | 0.9878 | 0.9048 |
| w2v | 2 | 870 | LR | 0.9529 | 0.9250 | 0.9742 | 0.8828 | 0.9879 | 0.9880 | 0.8943 |
| w2v | 5 | 1740 | LR | 0.9632 | 0.8783 | 0.9626 | 0.8103 | 0.9938 | 0.9860 | 0.8624 |
| LQL | 1 | 580 | LR | 0.9172 | 0.9116 | 0.9677 | 0.8655 | 0.9690 | 0.9849 | 0.8421 |
| LQL | 2 | 870 | LR | 0.9287 | 0.8821 | 0.9636 | 0.8172 | 0.9845 | 0.9819 | 0.8395 |
| LQL | 5 | 1740 | NB | 0.8316 | 0.4017 | 0.5192 | 0.3379 | 0.9303 | 0.7522 | 0.3226 |
| w2v | 1 | 580 | NB | 0.8897 | 0.8749 | 0.9879 | 0.7897 | 0.9897 | 0.9602 | 0.7976 |
| w2v | 2 | 870 | NB | 0.8529 | 0.7173 | 0.9891 | 0.5655 | 0.9966 | 0.9587 | 0.6743 |
| w2v | 5 | 1740 | NB | 0.9040 | 0.5845 | 1.0000 | 0.4241 | 1.0000 | 0.9577 | 0.6103 |
| LQL | 1 | 580 | NB | 0.6759 | 0.6300 | 0.7390 | 0.5552 | 0.7966 | 0.7684 | 0.3660 |
| LQL | 2 | 870 | NB | 0.7368 | 0.5483 | 0.6437 | 0.4828 | 0.8638 | 0.7419 | 0.3782 |
| LQL | 5 | 1740 | RF | 0.9644 | 0.8841 | 0.9616 | 0.8207 | 0.9931 | 0.9772 | 0.8680 |
| w2v | 1 | 580 | RF | 0.9414 | 0.9391 | 0.9706 | 0.9103 | 0.9724 | 0.9835 | 0.8850 |
| w2v | 2 | 870 | RF | 0.9540 | 0.9287 | 0.9647 | 0.8966 | 0.9828 | 0.9833 | 0.8968 |
| w2v | 5 | 1740 | RF | 0.9690 | 0.9009 | 0.9584 | 0.8517 | 0.9924 | 0.9806 | 0.8855 |
| LQL | 1 | 580 | RF | 0.9293 | 0.9273 | 0.9504 | 0.9069 | 0.9517 | 0.9781 | 0.8609 |
| LQL | 2 | 870 | RF | 0.9402 | 0.9052 | 0.9520 | 0.8655 | 0.9776 | 0.9765 | 0.8653 |
| LQL | 5 | 1740 | SVM | 0.9707 | 0.9076 | 0.9562 | 0.8655 | 0.9917 | 0.9825 | 0.8925 |
| w2v | 1 | 580 | KNN | 0.9534 | 0.9518 | 0.9718 | 0.9345 | 0.9724 | 0.9862 | 0.9091 |
| w2v | 1 | 580 | LR | 0.9500 | 0.9483 | 0.9682 | 0.9310 | 0.9690 | 0.9879 | 0.9021 |
| w2v | 5 | 1740 | SVM | 0.9701 | 0.9059 | 0.9340 | 0.8828 | 0.9876 | 0.9859 | 0.8899 |
| LQL | 1 | 580 | SVM | 0.9414 | 0.9387 | 0.9754 | 0.9069 | 0.9759 | 0.9894 | 0.8868 |
| LQL | 2 | 870 | SVM | 0.9448 | 0.9116 | 0.9581 | 0.8724 | 0.9810 | 0.9806 | 0.8753 |

The standard deviations of ACC(Accuracy), F1, RECALL, PRECISION, MCC, AUC, SPECIFICITY during 10-CV training were between 0.01-0.04

Table S2. related to Table 1

ALL models training results_2

| Protein codings | Ratio of negative/positive | Sequence number | Machine Learning algorithms | Accuracy | F1 | Precision | Recall | Specificty | AUC | MCC |
|---|---|---|---|---|---|---|---|---|---|---|
| LQL | 2 | 870 | DT | 0.9000 | 0.8524 | 0.8455 | 0.8621 | 0.9190 | 0.8903 | 0.7783 |
| w2v | 1 | 580 | DT | 0.9121 | 0.9097 | 0.9192 | 0.9034 | 0.9207 | 0.9121 | 0.8268 |
| LQL | 1 | 580 | DT | 0.8741 | 0.8752 | 0.8695 | 0.8828 | 0.8655 | 0.8741 | 0.7499 |
| w2v | 5 | 1740 | DT | 0.9534 | 0.8621 | 0.8505 | 0.8793 | 0.9683 | 0.9238 | 0.8362 |
| w2v | 2 | 870 | DT | 0.9115 | 0.8705 | 0.8543 | 0.8897 | 0.9224 | 0.9060 | 0.8050 |
| LQL | 5 | 1740 | DT | 0.9402 | 0.8191 | 0.8242 | 0.8172 | 0.9648 | 0.8926 | 0.7845 |
| w2v | 5 | 1740 | GBDT | 0.9770 | 0.9298 | 0.9483 | 0.9138 | 0.9897 | 0.9892 | 0.9169 |
| w2v | 1 | 580 | GBDT | 0.9638 | 0.9634 | 0.9662 | 0.9621 | 0.9655 | 0.9887 | 0.9289 |
| w2v | 2 | 870 | GBDT | 0.9632 | 0.9442 | 0.9501 | 0.9414 | 0.9741 | 0.9881 | 0.9184 |
| LQL | 1 | 580 | GBDT | 0.9362 | 0.9360 | 0.9417 | 0.9310 | 0.9414 | 0.9799 | 0.8731 |
| LQL | 2 | 870 | GBDT | 0.9598 | 0.9384 | 0.9544 | 0.9241 | 0.9776 | 0.9876 | 0.9094 |
| LQL | 5 | 1740 | GBDT | 0.9707 | 0.9083 | 0.9509 | 0.8724 | 0.9903 | 0.9861 | 0.8932 |
| LQL | 2 | 870 | KNN | 0.9322 | 0.8878 | 0.9746 | 0.8172 | 0.9897 | 0.9683 | 0.8476 |
| w2v | 1 | 580 | KNN | 0.9483 | 0.9476 | 0.9587 | 0.9379 | 0.9586 | 0.9834 | 0.8978 |
| LQL | 1 | 580 | KNN | 0.8914 | 0.8823 | 0.9480 | 0.8276 | 0.9552 | 0.9597 | 0.7905 |
| w2v | 5 | 1740 | KNN | 0.9701 | 0.9057 | 0.9530 | 0.8655 | 0.9910 | 0.9792 | 0.8905 |
| w2v | 2 | 870 | KNN | 0.9448 | 0.9136 | 0.9432 | 0.8897 | 0.9724 | 0.9777 | 0.8760 |
| LQL | 5 | 1740 | KNN | 0.9563 | 0.8508 | 0.9745 | 0.7586 | 0.9959 | 0.9617 | 0.8361 |
| LQL | 2 | 870 | LR | 0.9276 | 0.8801 | 0.9703 | 0.8069 | 0.9879 | 0.9776 | 0.8371 |
| LQL | 1 | 580 | LR | 0.9207 | 0.9160 | 0.9628 | 0.8759 | 0.9655 | 0.9762 | 0.8465 |
| w2v | 5 | 1740 | LR | 0.9632 | 0.8790 | 0.9528 | 0.8207 | 0.9917 | 0.9876 | 0.8627 |
| w2v | 2 | 870 | LR | 0.9563 | 0.9323 | 0.9572 | 0.9103 | 0.9793 | 0.9867 | 0.9016 |
| w2v | 1 | 580 | LR | 0.9500 | 0.9488 | 0.9686 | 0.9310 | 0.9690 | 0.9868 | 0.9018 |
| LQL | 5 | 1740 | LR | 0.9437 | 0.8017 | 0.9642 | 0.6897 | 0.9945 | 0.9785 | 0.7862 |
| LQL | 2 | 870 | NB | 0.7460 | 0.4882 | 0.7384 | 0.3690 | 0.9345 | 0.7826 | 0.3831 |
| w2v | 1 | 580 | NB | 0.8828 | 0.8670 | 0.9706 | 0.7897 | 0.9759 | 0.9614 | 0.7817 |
| LQL | 1 | 580 | NB | 0.6810 | 0.6298 | 0.7427 | 0.5517 | 0.8103 | 0.7523 | 0.3754 |
| w2v | 5 | 1740 | NB | 0.9023 | 0.5785 | 0.9762 | 0.4241 | 0.9979 | 0.9589 | 0.5995 |
| w2v | 2 | 870 | NB | 0.8575 | 0.7293 | 0.9804 | 0.5862 | 0.9931 | 0.9598 | 0.6834 |
| LQL | 5 | 1740 | NB | 0.8443 | 0.3005 | 0.5820 | 0.2103 | 0.9710 | 0.7771 | 0.2779 |
| LQL | 2 | 870 | RF | 0.9437 | 0.9098 | 0.9664 | 0.8621 | 0.9845 | 0.9794 | 0.8735 |
| w2v | 1 | 580 | RF | 0.9448 | 0.9432 | 0.9626 | 0.9276 | 0.9621 | 0.9788 | 0.8926 |
| LQL | 1 | 580 | RF | 0.9190 | 0.9163 | 0.9387 | 0.8966 | 0.9414 | 0.9760 | 0.8401 |
| w2v | 5 | 1740 | RF | 0.9701 | 0.9027 | 0.9720 | 0.8448 | 0.9952 | 0.9828 | 0.8891 |
| w2v | 2 | 870 | RF | 0.9540 | 0.9258 | 0.9705 | 0.8897 | 0.9862 | 0.9823 | 0.8968 |
| LQL | 5 | 1740 | RF | 0.9598 | 0.8686 | 0.9470 | 0.8069 | 0.9903 | 0.9811 | 0.8506 |
| LQL | 2 | 870 | SVM | 0.9517 | 0.9233 | 0.9694 | 0.8828 | 0.9862 | 0.9787 | 0.8911 |
| w2v | 5 | 1740 | SVM | 0.9741 | 0.9191 | 0.9533 | 0.8897 | 0.9910 | 0.9887 | 0.9054 |
| LQL | 1 | 580 | SVM | 0.9362 | 0.9337 | 0.9610 | 0.9103 | 0.9621 | 0.9807 | 0.8756 |
| w2v | 1 | 580 | SVM | 0.9603 | 0.9599 | 0.9659 | 0.9552 | 0.9655 | 0.9864 | 0.9220 |
| w2v | 2 | 870 | SVM | 0.9552 | 0.9320 | 0.9421 | 0.9241 | 0.9707 | 0.9835 | 0.8997 |
| LQL | 5 | 1740 | SVM | 0.9718 | 0.9105 | 0.9639 | 0.8655 | 0.9931 | 0.9777 | 0.8967 |

The standard deviations of ACC(Accuracy), F1, RECALL, PRECISION, MCC, AUC, SPECIFICITY during 10-CV training were between 0.01-0.04

Table S3. related to Table 1
ALL models training results_3

| Protein codings | Ratio of negative/positive | Sequence number | Machine Learning algorithms | Accuracy | F1 | Precision | Recall | Specificty | AUC | MCC |
|---|---|---|---|---|---|---|---|---|---|---|
| LQL | 5 | 1740 | DT | 0.9431 | 0.8303 | 0.8315 | 0.8310 | 0.9655 | 0.8983 | 0.7968 |
| w2v | 5 | 1740 | DT | 0.9523 | 0.8607 | 0.8407 | 0.8862 | 0.9655 | 0.9259 | 0.8340 |
| w2v | 2 | 870 | DT | 0.9103 | 0.8682 | 0.8592 | 0.8828 | 0.9241 | 0.9034 | 0.8034 |
| LQL | 2 | 870 | DT | 0.9046 | 0.8565 | 0.8602 | 0.8552 | 0.9293 | 0.8922 | 0.7864 |
| LQL | 1 | 580 | DT | 0.9069 | 0.9064 | 0.9018 | 0.9138 | 0.9000 | 0.9069 | 0.8162 |
| w2v | 1 | 580 | DT | 0.8914 | 0.8922 | 0.8841 | 0.9034 | 0.8793 | 0.8914 | 0.7858 |
| LQL | 5 | 1740 | GBDT | 0.9718 | 0.9119 | 0.9484 | 0.8793 | 0.9903 | 0.9866 | 0.8965 |
| w2v | 5 | 1740 | GBDT | 0.9770 | 0.9296 | 0.9479 | 0.9138 | 0.9897 | 0.9889 | 0.9167 |
| w2v | 2 | 870 | GBDT | 0.9632 | 0.9436 | 0.9618 | 0.9276 | 0.9810 | 0.9897 | 0.9175 |
| LQL | 2 | 870 | GBDT | 0.9494 | 0.9235 | 0.9331 | 0.9172 | 0.9655 | 0.9812 | 0.8876 |
| LQL | 1 | 580 | GBDT | 0.9621 | 0.9610 | 0.9724 | 0.9517 | 0.9724 | 0.9851 | 0.9258 |
| w2v | 1 | 580 | GBDT | 0.9621 | 0.9618 | 0.9639 | 0.9621 | 0.9621 | 0.9881 | 0.9262 |
| LQL | 5 | 1740 | KNN | 0.9603 | 0.8674 | 0.9750 | 0.7828 | 0.9959 | 0.9634 | 0.8521 |
| w2v | 5 | 1740 | KNN | 0.9730 | 0.9136 | 0.9777 | 0.8586 | 0.9959 | 0.9803 | 0.9008 |
| LQL | 2 | 870 | KNN | 0.9218 | 0.8688 | 0.9720 | 0.7897 | 0.9879 | 0.9489 | 0.8253 |
| LQL | 1 | 580 | KNN | 0.9052 | 0.8985 | 0.9606 | 0.8448 | 0.9655 | 0.9612 | 0.8168 |
| w2v | 1 | 580 | KNN | 0.9569 | 0.9559 | 0.9761 | 0.9379 | 0.9759 | 0.9860 | 0.9158 |
| w2v | 2 | 870 | KNN | 0.9563 | 0.9323 | 0.9645 | 0.9034 | 0.9828 | 0.9847 | 0.9019 |
| LQL | 5 | 1740 | LR | 0.9414 | 0.7924 | 0.9568 | 0.6793 | 0.9938 | 0.9783 | 0.7759 |
| w2v | 5 | 1740 | LR | 0.9655 | 0.8876 | 0.9614 | 0.8276 | 0.9931 | 0.9885 | 0.8721 |
| w2v | 2 | 870 | LR | 0.9540 | 0.9272 | 0.9741 | 0.8862 | 0.9879 | 0.9864 | 0.8967 |
| LQL | 2 | 870 | LR | 0.8977 | 0.8206 | 0.9653 | 0.7207 | 0.9862 | 0.9652 | 0.7710 |
| LQL | 1 | 580 | LR | 0.9345 | 0.9299 | 0.9768 | 0.8897 | 0.9793 | 0.9793 | 0.8739 |
| w2v | 1 | 580 | LR | 0.9552 | 0.9535 | 0.9787 | 0.9310 | 0.9793 | 0.9887 | 0.9126 |
| LQL | 5 | 1740 | NB | 0.8466 | 0.3079 | 0.6241 | 0.2069 | 0.9745 | 0.7765 | 0.2947 |
| w2v | 5 | 1740 | NB | 0.9029 | 0.5869 | 0.9864 | 0.4241 | 0.9986 | 0.9622 | 0.6077 |
| w2v | 2 | 870 | NB | 0.8609 | 0.7395 | 0.9764 | 0.5966 | 0.9931 | 0.9603 | 0.6900 |
| LQL | 2 | 870 | NB | 0.7276 | 0.4376 | 0.7019 | 0.3241 | 0.9293 | 0.7725 | 0.3306 |
| LQL | 1 | 580 | NB | 0.6879 | 0.6352 | 0.7591 | 0.5483 | 0.8276 | 0.7552 | 0.3913 |
| w2v | 1 | 580 | NB | 0.8897 | 0.8763 | 0.9722 | 0.8034 | 0.9759 | 0.9681 | 0.7943 |
| LQL | 5 | 1740 | RF | 0.9632 | 0.8784 | 0.9637 | 0.8103 | 0.9938 | 0.9816 | 0.8628 |
| w2v | 5 | 1740 | RF | 0.9690 | 0.9023 | 0.9447 | 0.8655 | 0.9897 | 0.9868 | 0.8859 |
| w2v | 2 | 870 | RF | 0.9460 | 0.9151 | 0.9527 | 0.8828 | 0.9776 | 0.9833 | 0.8783 |
| LQL | 2 | 870 | RF | 0.9448 | 0.9110 | 0.9739 | 0.8586 | 0.9879 | 0.9779 | 0.8765 |
| LQL | 1 | 580 | RF | 0.9310 | 0.9286 | 0.9539 | 0.9069 | 0.9552 | 0.9770 | 0.8649 |
| w2v | 1 | 580 | RF | 0.9483 | 0.9469 | 0.9692 | 0.9276 | 0.9690 | 0.9887 | 0.8991 |
| LQL | 5 | 1740 | SVM | 0.9678 | 0.8978 | 0.9549 | 0.8483 | 0.9917 | 0.9783 | 0.8814 |
| w2v | 5 | 1740 | SVM | 0.9753 | 0.9228 | 0.9637 | 0.8862 | 0.9931 | 0.9881 | 0.9096 |
| w2v | 1 | 580 | SVM | 0.9655 | 0.9645 | 0.9767 | 0.9552 | 0.9759 | 0.9874 | 0.9333 |
| w2v | 2 | 870 | SVM | 0.9621 | 0.9420 | 0.9622 | 0.9241 | 0.9810 | 0.9863 | 0.9151 |
| LQL | 2 | 870 | SVM | 0.9379 | 0.9006 | 0.9571 | 0.8552 | 0.9793 | 0.9694 | 0.8611 |
| LQL | 1 | 580 | SVM | 0.9362 | 0.9323 | 0.9741 | 0.8966 | 0.9759 | 0.9798 | 0.8769 |

The standard deviations of ACC(Accuracy), F1, RECALL, PRECISION, MCC, AUC, SPECIFICITY during 10-CV training were between 0.01-0.04

Table S4. related to main text
Enrichment GO terms of human PSPs predicted by PSPredictor

| GO terms | Percent(%)[a] | P-Value[b] |
|---|---|---|
| nucleus | 42.8 | 1.20E-53 |
| nucleoplasm | 25.4 | 8.10E-42 |
| cytoplasm | 35 | 2.00E-20 |
| nuclear speck | 3.6 | 4.80E-18 |
| collagen trimer | 2 | 1.40E-12 |
| intracellular ribonucleoprotein complex | 2.3 | 1.50E-11 |
| transcription factor complex | 2.7 | 4.10E-10 |
| nuclear chromatin | 2.6 | 1.40E-09 |
| postsynaptic density | 2.3 | 3.50E-07 |
| endoplasmic reticulum lumen | 2.3 | 1.00E-06 |
| intracellular membrane-bounded organelle | 4.7 | 2.30E-06 |
| actin cytoskeleton | 2.4 | 3.30E-06 |
| viral nucleocapsid | 0.7 | 4.60E-06 |
| nucleolus | 6.5 | 7.80E-06 |
| collagen type IV trimer | 0.4 | 1.80E-05 |
| nuclear pore central transport channel | 0.5 | 2.20E-05 |
| nuclear euchromatin | 0.7 | 2.30E-05 |
| lamellipodium | 1.8 | 5.80E-05 |
| cytoplasmic stress granule | 0.7 | 7.10E-05 |
| nuclear pore nuclear basket | 0.4 | 1.50E-04 |
| cytoskeleton | 3.1 | 1.60E-04 |
| messenger ribonucleoprotein complex | 0.3 | 5.20E-04 |
| cornified envelope | 0.7 | 7.50E-04 |
| proteinaceous extracellular matrix | 2.3 | 1.30E-03 |
| cell junction | 3.4 | 1.40E-03 |
| chromosome | 1.2 | 1.50E-03 |
| transcription elongation factor complex | 0.5 | 1.50E-03 |
| histone methyltransferase complex | 0.5 | 1.50E-03 |
| histone deacetylase complex | 0.6 | 1.60E-03 |
| Cajal body | 0.7 | 1.90E-03 |
| Z disc | 1.2 | 2.30E-03 |
| cytoplasmic mRNA processing body | 0.9 | 3.10E-03 |
| intermediate filament | 1.2 | 3.40E-03 |
| nBAF complex | 0.4 | 3.40E-03 |
| transcriptional repressor complex | 0.7 | 3.50E-03 |
| elongin complex | 0.3 | 3.60E-03 |
| synapse | 1.6 | 4.60E-03 |
| adherens junction | 0.7 | 5.30E-03 |
| catalytic step 2 spliceosome | 1 | 5.70E-03 |
| nuclear heterochromatin | 0.4 | 6.10E-03 |
| exon-exon junction complex | 0.4 | 6.10E-03 |
| bicellular tight junction | 1.1 | 7.70E-03 |
| nuclear transcription factor complex | 0.3 | 8.30E-03 |
| nuclear membrane | 1.8 | 8.30E-03 |
| BAF-type complex | 0.2 | 8.30E-03 |
| stress fiber | 0.7 | 9.20E-03 |
| nuclear matrix | 1 | 9.30E-03 |
| chromatin | 0.9 | 1.00E-02 |
| npBAF complex | 0.3 | 1.20E-02 |
| protein-DNA complex | 0.4 | 1.20E-02 |
| actin filament | 0.7 | 1.30E-02 |
| nuclear body | 0.5 | 1.30E-02 |
| PRC1 complex | 0.3 | 1.60E-02 |
| collagen type V trimer | 0.2 | 1.80E-02 |
| mRNA cleavage and polyadenylation specificity factor complex | 0.3 | 2.10E-02 |
| heterochromatin | 0.4 | 2.20E-02 |
| neuron projection | 1.8 | 2.20E-02 |
| filopodium | 0.7 | 2.40E-02 |
| dendritic spine | 0.9 | 2.60E-02 |
| SWI/SNF complex | 0.3 | 2.70E-02 |
| polysome | 0.5 | 2.80E-02 |
| clathrin-coated pit | 0.6 | 2.80E-02 |
| junctional sarcoplasmic reticulum membrane | 0.2 | 3.90E-02 |
| dendrite | 2.3 | 3.90E-02 |
| growth cone | 1 | 4.10E-02 |
| cytoplasmic ribonucleoprotein granule | 0.4 | 4.40E-02 |
| cytoplasmic, membrane-bounded vesicle | 1.1 | 4.60E-02 |
| basement membrane | 0.7 | 4.70E-02 |
| postsynaptic membrane | 1.5 | 4.70E-02 |
| perinuclear region of cytoplasm | 3.8 | 4.80E-02 |
| microtubule associated complex | 0.4 | 4.90E-02 |
| axonal growth cone | 0.3 | 4.90E-02 |
| nuclear inner membrane | 0.5 | 5.10E-02 |
| microtubule | 2.1 | 5.50E-02 |
| mediator complex | 0.4 | 5.50E-02 |
| cell-cell adherens junction | 2.2 | 5.60E-02 |
| PcG protein complex | 0.4 | 5.80E-02 |
| paraspeckles | 0.2 | 7.60E-02 |
| CRD-mediated mRNA stability complex | 0.2 | 7.60E-02 |
| clathrin-coated vesicle | 0.6 | 8.60E-02 |
| ruffle | 0.7 | 9.90E-02 |

a. Percent is the ratio of the number of proteins that were annotated with the GO terms and the total number of proteins
b. p-value is a modified Fisher Exact P-Value, for gene-enrichment analysis. It ranges from 0 to 1. Fisher Exact P-Value = 0 represents perfect enrichment. Usually P-Value is equal or smaller than 0.05 to be considered strongly enriched in the annotation

Table S5. related to main text

GO term clusters of PSPs with similar meaning in biology

| | | | | |
|---|---|---|---|---|
| Annotation ( | Enrichment Score: 7.94 | | Count | P_Value |
| | GOTERM_CC_DIRECT | collagen trimer | 32 | 1.40E-12 |
| | GOTERM_MF_DIRECT | extracellular matrix structural constituent | 26 | 3.10E-11 |
| | GOTERM_BP_DIRECT | collagen catabolic process | 25 | 4.60E-11 |
| | GOTERM_BP_DIRECT | extracellular matrix organization | 45 | 2.80E-10 |
| | GOTERM_CC_DIRECT | endoplasmic reticulum lumen | 37 | 1.00E-06 |
| | GOTERM_BP_DIRECT | collagen fibril organization | 13 | 3.50E-05 |
| | GOTERM_CC_DIRECT | proteinaceous extracellular matrix | 37 | 1.30E-03 |
| Annotation ( | Enrichment Score: 3.15 | | Count | P_Value |
| | GOTERM_MF_DIRECT | translation repressor activity, nucleic acid binding | 8 | 2.30E-06 |
| | GOTERM_BP_DIRECT | negative regulation of cytoplasmic translation | 6 | 6.10E-05 |
| | GOTERM_CC_DIRECT | messenger ribonucleoprotein complex | 5 | 5.20E-04 |
| | GOTERM_MF_DIRECT | mRNA 3'-UTR AU-rich region binding | 6 | 1.10E-03 |
| | GOTERM_MF_DIRECT | translation factor activity, RNA binding | 7 | 2.80E-02 |
| | GOTERM_MF_DIRECT | ribosome binding | 9 | 5.40E-02 |
| Annotation ( | Enrichment Score: 2.99 | | Count | P_Value |
| | GOTERM_BP_DIRECT | RNA export from nucleus | 16 | 1.90E-05 |
| | GOTERM_BP_DIRECT | mRNA export from nucleus | 20 | 3.40E-04 |
| | GOTERM_BP_DIRECT | mRNA 3'-end processing | 11 | 5.70E-03 |
| | GOTERM_BP_DIRECT | termination of RNA polymerase II transcription | 11 | 3.10E-02 |
| Annotation ( | Enrichment Score: 2.85 | | Count | P_Value |
| | GOTERM_BP_DIRECT | RNA export from nucleus | 16 | 1.90E-05 |
| | GOTERM_CC_DIRECT | nuclear pore central transport channel | 8 | 2.20E-05 |
| | GOTERM_CC_DIRECT | nuclear pore nuclear basket | 7 | 1.50E-04 |
| | GOTERM_MF_DIRECT | structural constituent of nuclear pore | 8 | 1.00E-03 |
| | GOTERM_MF_DIRECT | nucleocytoplasmic transporter activity | 8 | 1.40E-03 |
| | GOTERM_BP_DIRECT | protein sumoylation | 20 | 2.40E-03 |
| | GOTERM_BP_DIRECT | protein import into nucleus | 12 | 5.80E-03 |
| | GOTERM_MF_DIRECT | nuclear localization sequence binding | 8 | 6.20E-03 |
| | GOTERM_BP_DIRECT | regulation of cellular response to heat | 14 | 6.30E-03 |
| | GOTERM_BP_DIRECT | mitotic nuclear envelope disassembly | 10 | 7.30E-03 |
| | GOTERM_BP_DIRECT | tRNA export from nucleus | 8 | 1.20E-02 |
| | GOTERM_BP_DIRECT | regulation of glucose transport | 8 | 1.40E-02 |
| Annotation ( | Enrichment Score: 2.7 | | Count | P_Value |
| | GOTERM_BP_DIRECT | keratinocyte differentiation | 17 | 2.90E-04 |
| | GOTERM_CC_DIRECT | cornified envelope | 12 | 7.50E-04 |
| | GOTERM_BP_DIRECT | keratinization | 11 | 4.20E-03 |
| | GOTERM_BP_DIRECT | peptide cross-linking | 10 | 1.70E-02 |
| Annotation ( | Enrichment Score: 2.09 | | Count | P_Value |
| | GOTERM_CC_DIRECT | histone methyltransferase complex | 8 | 1.50E-03 |
| | GOTERM_MF_DIRECT | histone-lysine N-methyltransferase activity | 9 | 1.20E-02 |
| | GOTERM_MF_DIRECT | histone methyltransferase activity (H3-K4 specific) | 6 | 1.30E-02 |
| | GOTERM_BP_DIRECT | histone H3-K4 methylation | 6 | 1.90E-02 |
| Annotation ( | Enrichment Score: 2.03 | | Count | P_Value |
| | GOTERM_CC_DIRECT | nBAF complex | 6 | 3.40E-03 |
| | GOTERM_BP_DIRECT | ATP-dependent chromatin remodeling | 7 | 8.10E-03 |
| | GOTERM_CC_DIRECT | BAF-type complex | 4 | 8.30E-03 |
| | GOTERM_CC_DIRECT | npBAF complex | 5 | 1.20E-02 |
| | GOTERM_CC_DIRECT | SWI/SNF complex | 5 | 2.70E-02 |
| Annotation ( | Enrichment Score: 2.02 | | Count | P_Value |
| | GOTERM_BP_DIRECT | nucleosome positioning | 5 | 2.20E-03 |
| | GOTERM_BP_DIRECT | histone H3-K4 trimethylation | 6 | 5.00E-03 |
| | GOTERM_BP_DIRECT | histone H3-K27 trimethylation | 3 | 7.80E-02 |
| Annotation ( | Enrichment Score: 1.89 | | Count | P_Value |
| | GOTERM_BP_DIRECT | positive regulation of excitatory postsynaptic potential | 7 | 3.80E-03 |
| | GOTERM_BP_DIRECT | protein localization to synapse | 5 | 1.60E-02 |
| | GOTERM_BP_DIRECT | regulation of alpha-amino-3-hydroxy-5-methyl-4-isoxazole propionate selective glutamate receptor activity | 5 | 3.40E-02 |
| Annotation ( | Enrichment Score: 1.64 | | Count | P_Value |
| | GOTERM_BP_DIRECT | positive regulation of excitatory postsynaptic potential | 7 | 3.80E-03 |
| | GOTERM_BP_DIRECT | vocalization behavior | 5 | 2.20E-02 |
| | GOTERM_BP_DIRECT | NMDA glutamate receptor clustering | 3 | 5.50E-02 |
| | GOTERM_BP_DIRECT | positive regulation of synaptic transmission, glutamatergic | 5 | 6.10E-02 |
| Annotation ( | Enrichment Score: 1.53 | | Count | P_Value |
| | GOTERM_BP_DIRECT | social behavior | 10 | 1.30E-02 |
| | GOTERM_BP_DIRECT | vocal learning | 4 | 1.40E-02 |
| | GOTERM_BP_DIRECT | vocalization behavior | 5 | 2.20E-02 |
| | GOTERM_MF_DIRECT | GKAP/Homer scaffold activity | 3 | 3.60E-02 |
| | GOTERM_BP_DIRECT | adult behavior | 6 | 5.30E-02 |
| | GOTERM_MF_DIRECT | ionotropic glutamate receptor binding | 5 | 8.70E-02 |
| Annotation ( | Enrichment Score: 1.24 | | Count | P_Value |
| | GOTERM_BP_DIRECT | cell-cell adhesion | 31 | 4.50E-02 |
| | GOTERM_CC_DIRECT | cell-cell adherens junction | 35 | 5.60E-02 |
| | GOTERM_MF_DIRECT | cadherin binding involved in cell-cell adhesion | 32 | 7.60E-02 |

Figure S1. Training and Test dataset construction. The number of overall conditions associated entries, protein sequences and proteins in LLPSDB beta

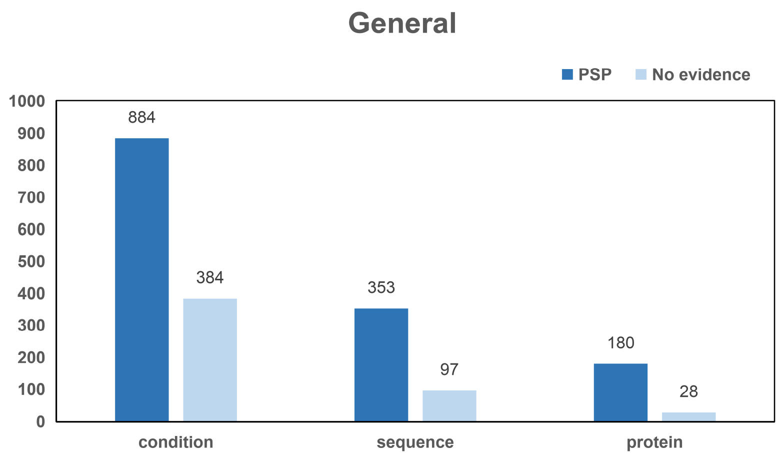
